# Progressive myoclonic epilepsy-associated gene *Kctd7* regulates retinal neurovascular patterning and function

**DOI:** 10.1101/647008

**Authors:** Jonathan Alevy, Courtney A. Burger, Nicholas E. Albrecht, Danye Jiang, Melanie A. Samuel

**Affiliations:** Department of Neuroscience, Baylor College of Medicine, Houston, TX 77030; Huffington Center on Aging, Baylor College of Medicine, Houston, TX 77030

## Abstract

Neuron function relies on and instructs the development and precise organization of neurovascular units that in turn support circuit activity. However, our understanding of the molecular cues that regulate this relationship remains sparse. Using a high-throughput screening pipeline, we recently identified several new regulators of vascular patterning. Among these was the potassium channel tetramerization domain-containing protein 7 (KCTD7). Mutations in *KCTD7* are associated with progressive myoclonic epilepsy, but how KCTD7 regulates neural development and function remains poorly understood. To begin to identify such mechanisms, we focus on mouse retina, a tractable part of the central nervous system that contains precisely ordered neuron subtypes supported by a trilaminar intravascular network. We find that deletion of *Kctd7* results in defective patterning of the adult retina vascular network, resulting in increased branching, vessel length, and lacunarity. These alterations reflect early and specific defects in vessel development, as emergence of the superficial and deep vascular layers were delayed. These defects are likely due to a role for Kctd7 in inner retina neurons. Kctd7 it is absent from vessels but present in neurons in the inner retina, and its deletion resulted in a corresponding increase in the number of bipolar cells in development and increased vessel branching in adults. These alterations were accompanied by retinal function deficits. Together, these data suggest that neuronal Kctd7 drives growth and patterning of the vasculature and suggest that neurovascular interactions may participate in the pathogenesis of KCTD7-related human diseases.

**Alevy et al. Highlights:**

- Kctd7 is required for normal retinal vascular organization and retinal function in adults.
- Deletion of *Kctd7* disrupts the emergence of the superficial and deep vessel layers.
- Kctd7 may impact vascular patterning through influencing neurons as it is expressed in and regulates bipolar cells.
- Kctd7 driven neurovascular interactions may participate in the pathogenesis of KCTD7-related human diseases.

**Lay Summary:** Neuron function requires an organized vasculature to maintain brain health and prevent disease, but many neurovasculature regulatory genes remain unknown. Alevy et al. identify the progressive myoclonic epilepsy-associated gene *Kctd7* as a key regulator of vascular development and retinal function. They further show that Kctd7 regulation of vessel patterning likely occurs downstream of its role in regulating the development or activity of specific neuron types. These data suggest that KCTD7-regulated neurovascular interactions may participate in the pathogenesis of associated human diseases.

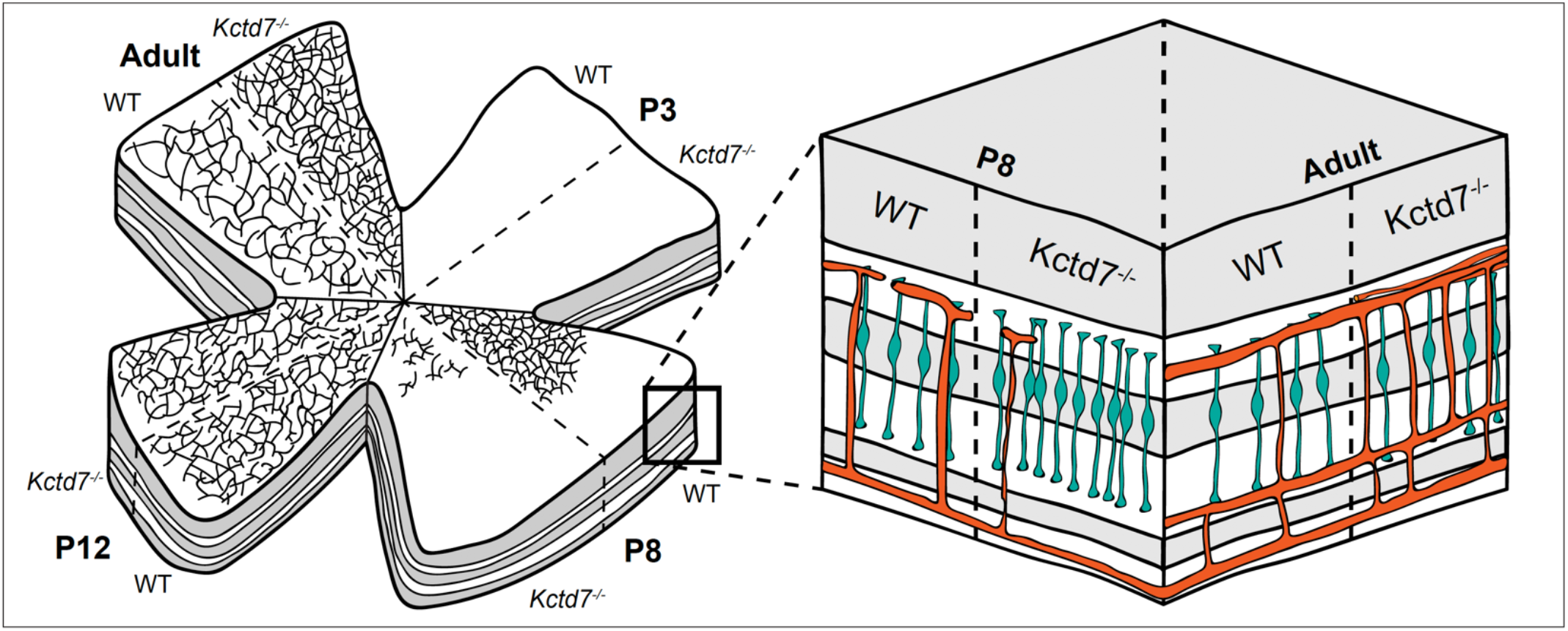

## 1. Introduction

The central nervous system (CNS) is one of the most energetically demanding parts of the body, and it relies on vasculature to appropriately meet the metabolic demand of neuron signaling, circuit function, and ultimately behavior. Correspondingly, a large number of neural diseases are associated with vascular defects, including those that are the most common and the most costly (e.g. Alzheimer’s disease and Parkinson’s disease, reviewed in Kisler et al., 2017; Sweeny et al., 2018). Yet, we know relatively little about the spectrum of molecules that can instruct neurovascular interactions or mediate pathogenic or protective outcomes in disease. In part, this is because the CNS is made up of thousands of types of neurons, each of which have unique properties, signaling modalities, and energetic demands. Since these cell types and circuits are only beginning to be mapped, it has remained difficult to decode the contribution of neuron subtypes and molecules to vascular outcomes.

To overcome these challenges, we focus here on the murine retina. Retina neurons are ordered in precise layers termed lamina that form an organized array of distinct neuron subsets (approximately ~100 types, Macosko et al., 2015) which can be mapped and manipulated. Signaling in this circuit begins with photoreceptors, which reside in the outer nuclear layer (ONL) and detect light. Visual signals are then relayed through synapses located in the outer plexiform layer (OPL) to bipolar cells, amacrine cells, and horizontal cells in the inner nuclear layer (INL). Bipolar cells synapse in the inner plexiform layer (IPL) with retinal ganglion cells (RGCs) in the ganglion cell layer (GCL). RGCs then transmit visual information to the brain (Dowling, 2012).

In mouse, as in human, retinal neurons are supported by a trilaminar intraretinal vascular network (Stahl et al., 2010; Usui et al., 2015a). The vascular layers consist of the superficial plexus, which interdigitates the GCL, the intermediate plexus, which ascends into the IPL, and the deep plexus which interleaves the OPL (Stahl et al., 2010). These vessel layers function as neurovascular units comprised of endothelial cells, mural cells, astrocytes, and neurons (Rhodes and Simons, 2007; Ladecola, 2017). In turn, each layer has a characteristic location, branching pattern, and complexity. Because the pattern of these networks is stereotyped and influenced by local neuron interactions (Usui et al., 2015b; Selvam et al., 2017; Paredes et al., 2018), the retina vascular layers provide an ideal system to dissect neurovascular interactions.

Given these features, we recently established a pipeline for using mouse retina to identify new regulators of neurons, vasculature, and their interactions (Albrecht et al., 2018). This screen implicated several novel genes in retinal vasculature organization, including the potassium channel tetramerization domain-containing protein 7 (*Kctd7*). This gene is of particular interest for two reasons. First, it is implicated in progressive myoclonic epilepsy, a rare but serious human neural disease characterized by the presence of abnormal muscle contraction and seizures (Azizieh et al., 2011; Kousi et al., 2012; Staropoli et al., 2012; Van Bogaert, 2016). Since mouse models for this disease have not been available, its pathogenesis is not well understood. Second, KCTD7 is part of a large family of related proteins that regulate a diverse array of functions ranging from potassium channel tetramerization to ubiquitination (Kreusch et al. 1998; Krek, W, 2003; Genschik et al., 2013; Liu et al., 2013). But whether and how such proteins can influence the vasculature remains unknown.

Here, we show that Kctd7 is a regulator of vasculature organization and development in specific regions of the retina. Loss of Kctd7 resulted in increased vessel branching and vessel area in the deep and intermediate vessel layers in adults. Further, emergence of the superficial and deep vasculature layer was delayed, and deep layer *Kctd7*^−/−^ vessels were less complex early in development. These alterations likely reflect a neuron-specific role for *Kctd7* as its expression appears restricted to neurons. Correspondingly, alterations to the deep vascular layer were accompanied by a Kctd7-dependent increase in bipolar cell numbers in the first postnatal week. These retinal defects were accompanied by declines in retinal function. These studies are the first to implicate Kctd7 in vasculature regulation and suggest that neuron driven vascular changes could influence KCTD7-related human neurological diseases.

## 2. Materials and Methods

### 2.1. Mouse Strains

The *Kctd7*-null mutant (*Kctd7*^−/−^) was was provided by the International Mouse Phenotyping Consortium and was generated on a C57BL/6NJ background. Age matched, wildtype C57BL/6NJ controls were used for all studies and approximately equal numbers of male and female mice were included. Experiments were carried out in accordance with the Guide for the Care and Use of Laboratory Animals of the NIH under protocols approved by the BCM Institutional Animal Care and Use Committee (IACUC).

### 2.2. Tissue preparation and immunohistochemistry

Eyes were collected from wildtype and *Kctd7*^−/−^ animals at P3, P8, P12, and 16-weeks and fixed for 45 minutes in 4% paraformaldehyde (w/v) in phosphate buffered saline (PBS) and rinsed with PBS. Antibody information, dilutions, and specificity are detailed in **Supplemental Table 1**. For cross-section analysis, the optic cup was cryoprotected in 30% sucrose, embedded in Optimal Cutting Temperature (OCT) compound (VWR), frozen in methyl butane on dry ice, sectioned at 20 μm, and then mounted on Superfrost slides. Slides were incubated in blocking solution (3% normal donkey serum and 0.3% Triton X-100 [vol/vol] in PBS) for 1 hour, followed by incubation with primary antibodies overnight at 4°C and secondary antibodies for 1 hour at room temperature. For whole mount staining of the vasculature, the retina was removed from the optic cup following fixation and then incubated in blocking solution (10% normal donkey serum and 0.5% Triton X-100 [vol/vol] in PBS) followed by primary antibody labeling at 4°C for 7 days and secondary antibody labeling at 4°C for an additional 7 days. For vascular cross-section staining, fixed retinas were removed from the optic cup and embedded in 5% low melting point agarose. A vibratome (Leica VT1200) was used to generate 100 μm sections, which were then incubated in blocking solution (10% normal donkey serum and 0.5% Triton X-100 [vol/vol] in PBS) followed by primary antibody labeling at 4°C for 3 days and secondary antibody labeling at 4°C for an additional 3 days. All samples were mounted in Vectashield (Vectorlabs). Images were acquired on an Olympus FluoView FV1200 confocal microscope and processed using Fiji. Images were acquired with the following specifications: antibody cross section, 211 μm × 211 μm; whole mount, 635 μm × 635 μm; vessel cross section, 211 μm × 211 μm; and *in situ*, 211μm × 211 μm.

### 2.3 Histological quantification

For quantification, images were collected from 5 to 10 animals per group with at least 3 image stacks per animal. For vasculature quantification, retinas were harvested and processed for whole mount staining using an antibody to CD31. Images were acquired at equivalent retinal eccentricities from the optic nerve head and processed using Fiji. The number of branch points were quantified in merged images from 20-50 consecutive optical sections in order to separately visualize the superficial, intermediate, and deep layers. To quantify the total vessel area and mean lacunarity, we employed automated segmentation and reconstruction using Angiotool v0.6a. (Zudaire et al., 2011). To determine the relative retinal area occupied by vessel layers over development, retinas were whole mounted, and the area covered by the vasculature was computed relative to the total retina area. To examine cones, bipolars, amacrines, horizontal cells, and retinal ganglion cells and quantify their numbers, we used antibodies that mark each cell type specifically (**Supplemental Table 1**). Two image stacks were acquired at equivalent retinal eccentricities from the optic nerve head per animal. The total number of soma for each cell type was quantified in a standardized 211μm × 211μm optical sections from 5-10 non-successive optical sections per image stack.

### 2.4 In situ hybridization

*In situ* hybridization was performed by the RNA *In Situ* Hybridization Core at BCM using an automated robotic platform as previously described (Yaylaoglu et al., 2005). We prepared digoxigenin (DIG)- labeled riboprobes using reverse-transcribed mouse cDNA as a template that was generated from RNA harvested from mouse brain at E15 and P7. First strand cDNA synthesis was performed using the Superscript IV First-Strant Synthesis System (Invitrogen). Polymerase chain reaction (PCR) primers were used to generate cDNA fragments corresponding to the desired riboprobes for *Cspg4* and *Kctd7* (**Supplemental Table 2**). DIG- labeled riboprobes were synthesized using a DIG RNA labeling kit (Roche) and stored in hybridization buffer at a concentration of 100 ng/μl at stored at −20°C.

For *in situ* hybridization retinas were cryoprotected in 30% sucrose, frozen in OCT (VWR), cryosectioned 20μm, and mounted on Superfrost Plus slides (VWR). Sections were fixed and acetylated before the hybridization procedure, which was performed on a high-throughput platform. The slides were developed using tyramide labeled with Cy3 directly (TSA-Plus system; Perkin-Elmer Life Sciences) for 15 min and then stained with 4’,6-diamidino-2-phenylindole (DAPI) before mounting in Prolong Diamond (Invitrogen).

### 2.5 ERG

We performed ERG on adult 16 week-old animals controls as previously described (Albrecht *et al.*, 2018). In brief, mice (n = 7 mutants and 8 wildtype) were dark adapted overnight and anesthetized with 1.5% isoflurane at an oxygen flow rate of 1.0 L/min. Mice were placed on a heated platform and phenylephrine hydrochloride and tropicamide was used to topically dilate the pupil. Electroretinograms were monitored from both eyes simultaneously with a contact lens-style electrode in contact with Gonak solution on each cornea. Scotopic responses were elicited in the dark with flashes ranging from 0.003 cd*s/m^2^ to 20.0 cd*s/m^2^, and photopic responses were elicited with flashes at 0.003 cd*s/m^2^ and 10.0 cd*s/m^2^ using the Diagnosys Celeris ERG system. The ground electrode was subcutaneously placed into the forehead, and a reference electrode was placed into the hip.

### 2.7 Statistical Analysis

Statistical analyses on neuron numbers and ERG results were performed using an unpaired, two-tailed Student’s *t* test. Vascular branch number, total area, lacunarity, and quantification data were analyzed using a two-way ANOVA using the animal as the unit of analysis and allowing for unequal variance. No statistical analysis was conducted to pre-determine sample sizes. Randomization and blinding were not employed. Statistical differences were evaluated using Graphpad Prism 7 software. P<0.05 was considered statistically significant.

## 3. Results

### 3.1 Kctd7 is required for vascular organization

To examine the role of Kctd7 in retina vascular organization, we obtained *Kctd7*-null animals from the International Mouse Phenotyping Consortium (IMPC, Koscielny et al., 2014). *Kctd7*^−/−^ mice are viable and fertile (http://www.mousephenotype.org/), which allowed us to examine the role of Kctd7 in adults. To visualize the vasculature, we stained for CD31, which labels platelet-endothelial cell adhesion molecule 1 and can be used to visualize vessel structure (**Fig. 1A)**. Wildtype retinas from 16-week-old animals showed characteristic and distinctive segregation of the superficial, intermediate, and deep layers that displayed normal branching complexity and total vessel area (133 ± 24, 175 ± 58, 311 ± 51 branch points/mm^2^ in the superficial, intermediate, and deep layers, respectively, **Fig. 1A-B**). Like control animals, *Kctd7*^−/−^ mice displayed grossly normal organization of the superficial vascular layer (146 ± 21 branch points/mm^2^, P=0.92). However, both the intermediate and deep layers displayed a significant increase in vessel branching, resulting in ~30% more vessel branches. (271 ± 38 and 498 ± 62 /mm^2^ in the intermediate, and deep layers, respectively, P<0.001, **Fig. 1A-B**). The total area covered by the vasculature in the intermediate and deep layers was also correspondingly increased (54% and 22% increase, respectively, P<0.001, **Fig. 1C)**. To examine whether these changes impacted the degree of vessel homogeneity we performed a lacunarity analysis. Lacunarity was significantly reduced in *Kctd7*^−/−^ mice in both the deep and intermediate layers, with the intermediate layer being the most dramatically impacted (33% reduction and 49% reduction, respectively, P<0.001, **Fig. 1D**). Together, these results indicate that Kctd7 is required for patterning the intermediate and deep vasculature layers in adults.

**Figure 1.**
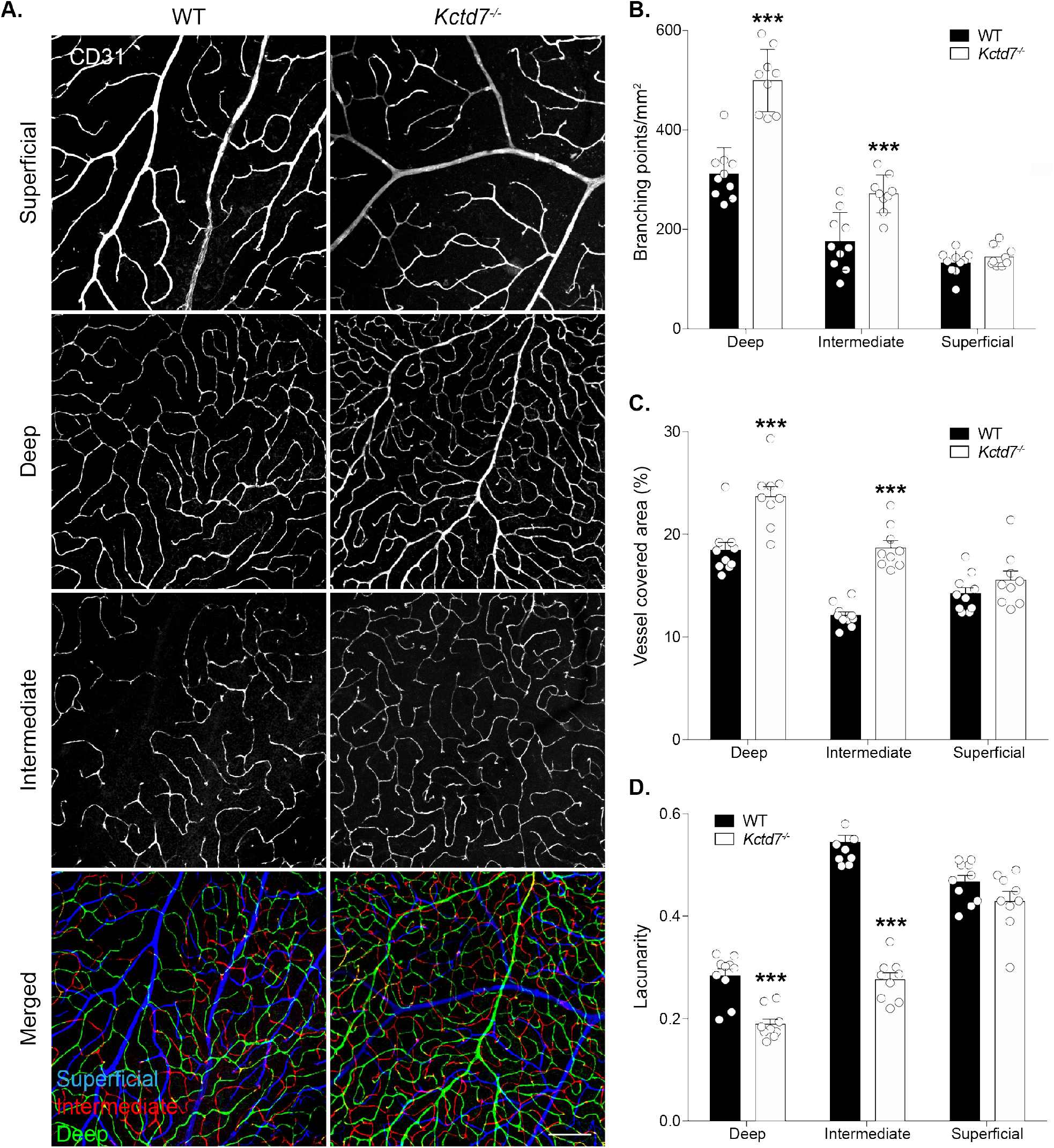
Vascular organization defects in *Kctd7*^−/−^ mice. **A.** To examine retina vascularization, 16-week-old wildtype control and *Kctd7*^−/−^ retinas were stained with an antibody to CD31. Representative images of the deep, intermediate, and superficial vascular layers are shown. Noted increases in vascular branching were observed in the intermediate and deep vessel layers. **B**. Differences in vessel branching were quantified in wildtype controls and *Kctd7*^−/−^ mice in each of the vascular layers by counting the number of vascular branch points following staining with CD31. Significant alterations were observed in the deep and intermediate layers. **C-D**. *Kctd7*^−/−^ intermediate and deep vessel layers showed an increase in the mean vessel covered area (**C**) and a decrease in lacunarity (**D**) relative to wildtype mice following automated segmentation, reconstruction, and quantification using Angiotool v0.6. n=10 wildtype and 9 *Kctd7*^−/−^ animals. Data are represented as the mean ± the s.e.m. Scale bars = 100μm. *** P<0.001, ** P<0.01, * P<0.05, 2way ANOVA and Dunnett’s multiple comparison test for significance.

### 3.2. Intraretinal vasculature layer development is disrupted in Kctd7 mutant mice

To determine if defects in adult vasculature reflected alterations in retinal vasculature development, we examined the retina at early postnatal ages when the vascular layers emerge (Dorrell, Friedlander, 2006; Stahl et al., 2010). Under normal conditions, intraretinal vascular layers originate when endothelial cells migrate from the optic nerve onto the retinal surface forming the superficial plexus in a process that begins at postnatal day (P) 0. Superficial capillaries sprout vertically from the superficial layer into the OPL, and vessels then extend laterally to form the deep plexus from P8 to P12. Beginning at P12, vessels from the deep plexus ascend into the IPL to form the intermediate plexus. Vascular development is complete by P15 (**Fig. 2A**). To examine this process, we first visualized vasculature layer restriction in retinal cross sections. Restriction of the superficial, deep, and intermediate vessel layers to their respective retinal layers was largely intact in *Kctd7*^−/−^ mice (**Supplemental Figure 1).** We next visualized and reconstructed individual vessel layers in whole mount preparations to quantify their emergence and spread across the retina over time. Like control animals, *Kctd7*^−/−^ mice displayed normal development of the superficial layer at P3 (16% versus 17% of the retinal area covered by vessels in wildtype and *Kctd7*^−/−^ mice respectively, **Fig. 2B-D**). However, as the superficial layer grew, defects in *Kctd7*^−/−^ mice emerged. While wildtype animals displayed a superficial plexus that covered the majority of the retina at P8 (92%), *Kctd7*^−/−^ mice showed a small but significant reduction in superficial layer growth, averaging 84% coverage at this time point (P<0.01, **Fig. 2D**). In parallel, branching of the superficial plexus was increased in *Kctd7*^−/−^ mice relative to wildtype controls at P8 (501 ± 129 versus 350 ± 52 branch points/mm^2^ respectively, P<0.01, **Fig. 2E**).

**Figure 2:**
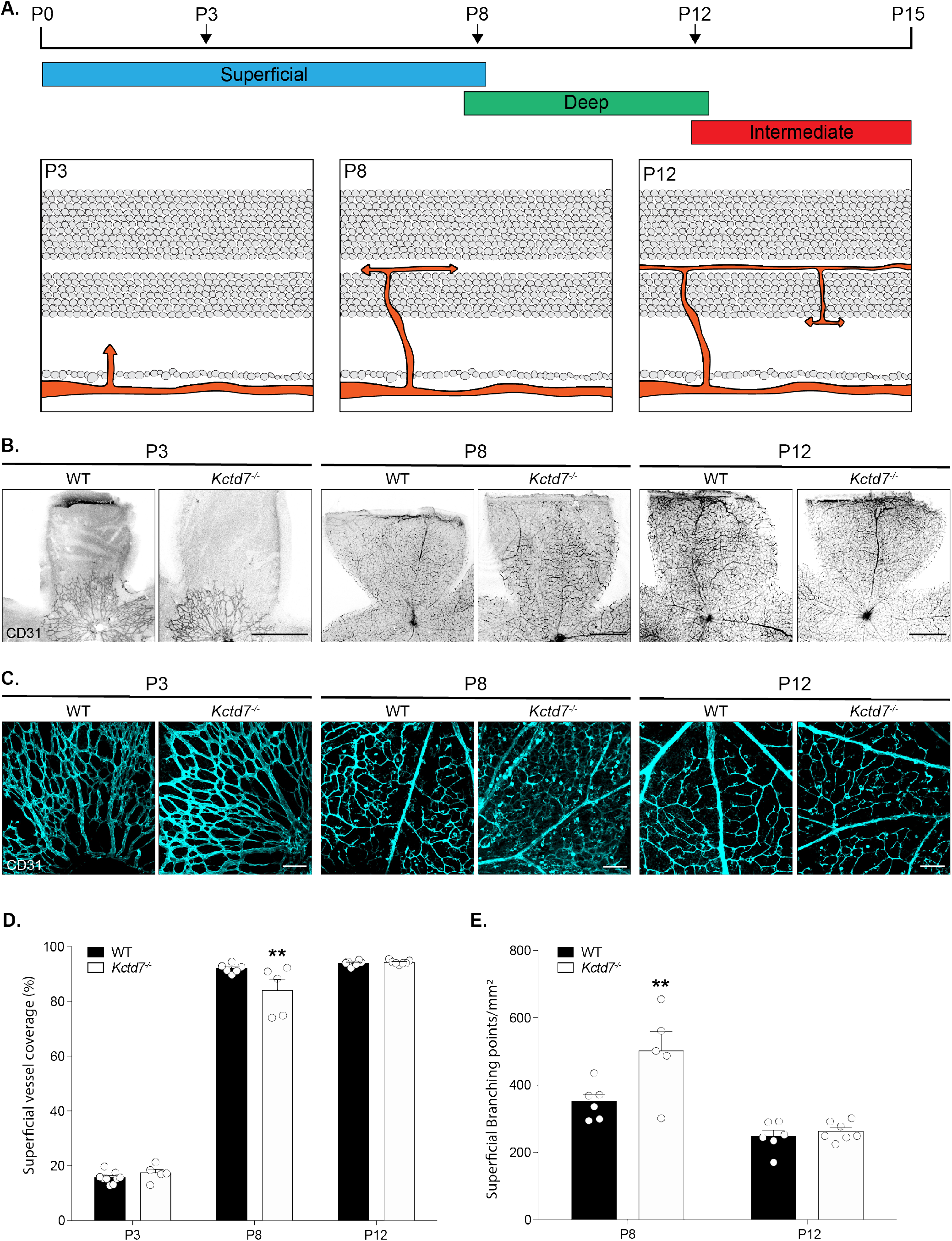
Kctd7 modulates the development of the superficial vascular plexus. **A.** Schematic of vasculature development from P0 to P15. The primary superficial plexus begins to emerge at P0, grows laterally to the periphery, and reaches its full radial growth by P8. From day 7, capillaries ascend into the OPL to form the deep plexus, which then interdigitates this layer and is completed by P12. Capillaries from the deep layer then descend into the IPL to form the intermediate layer between P12 and P15. By P15 all three layers are present. **B-C**. To visualize vascular development, whole mount retinas from wildtype and *Kctd7*^−/−^ animals were stained for CD31 at P3, P8, and P12. In wildtype mice, radial development of the superficial layer is apparent at P3 and complete by P8. Vessels grow in complexity until P12 (**B**, scale bars = 250μm), forming elaborate branching patterns (**C**, scale bars = 100μm). In *Kctd7*^−/−^ animals, development of the nascent superficial layer is intact at P3 (**B-C**). At P8, however, *Kctd7*^−/−^ animals showed a significant reduction in blood vessel coverage but an increase in branching complexity (**D**, **E).** n ≥ 6 wildtype and ≥ 5 *Kctd7*^−/−^ animals. Data are represented as the mean ± the s.e.m. *** P<0.001, ** P<0.01, * P<0.05, 2way ANOVA and Dunnett’s multiple comparison test for significance.

Since the development of the superficial layer at P8 coincides with the emergence of the deep vasculature layer, we examined whether these events impacted deep layer formation. To begin this analysis, we built on previous work to describe the time course in which the deep layer emerges and then covers the retina surface. In wildtype mice, the deep layer was not present at P3, but it could be readily detected at P8 when it occupied 53% of the retinal surface (**Fig. 3A-C**). In contrast, emergence and extension of the deep vessel layer was reduced in *Kctd7*^−/−^ mice, resulting in a 43% reduction in coverage at this time point relative to wildtype mice (P<0.001, **Fig.3A-C**). In addition, regions that were occupied by the deep vascular plexus showed a dramatically reduced vessel density, resulting in an 84% reduction in deep layer vessel branching in *Kctd7*^−/−^ mice (P<0.001; **Fig.3B, D**). The alterations in vascular development persisted until P12, when the intermediate layer began to emerge **(Fig. 3B-D**). Together, these data indicate that Kctd7 is required for proper temporal emergence and patterning of the intraretinal vasculature and particularly impacts the deep vascular layer.

**Figure 3:**
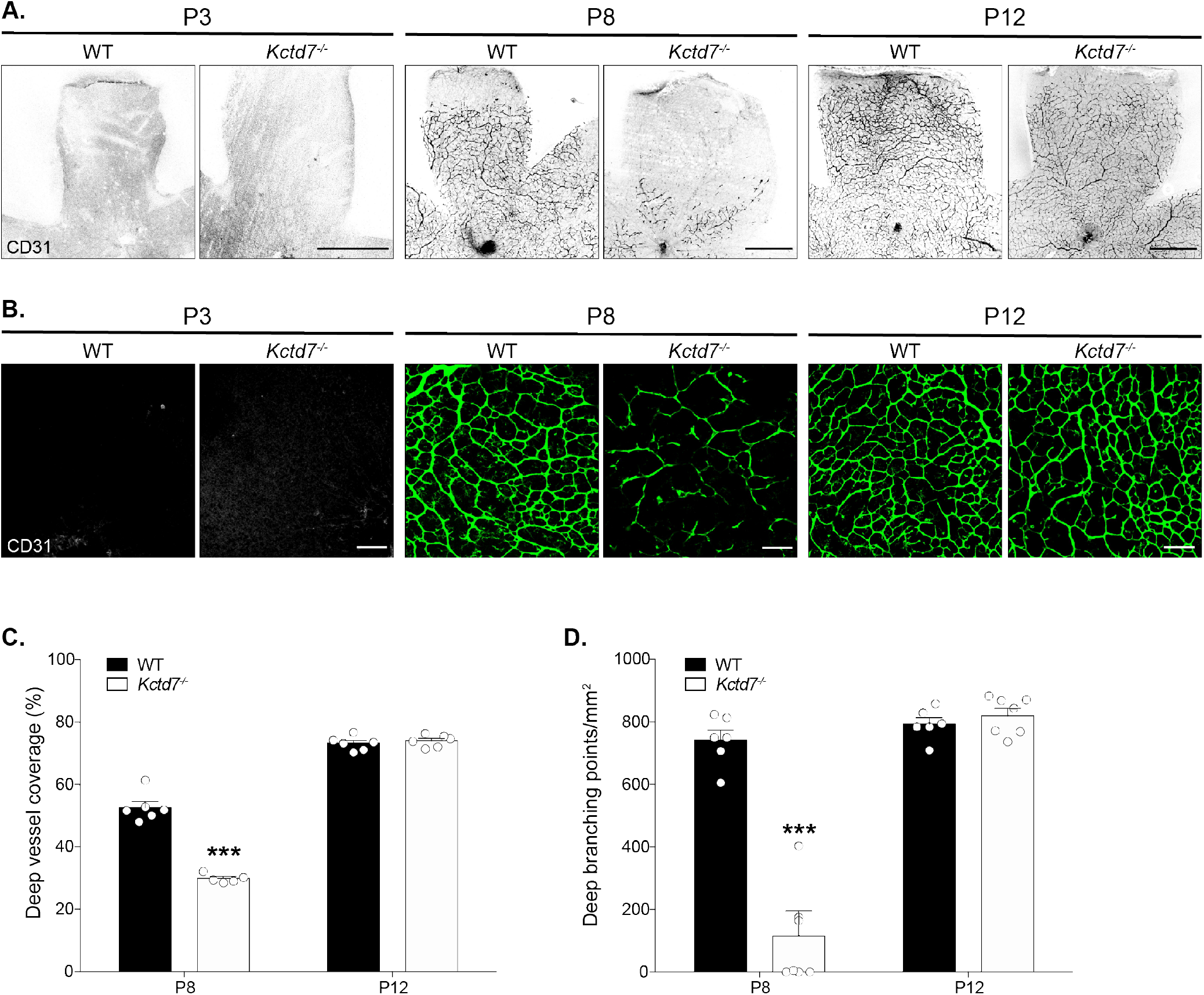
Kctd7 is required for the emergence of the deep vascular layer. **A-B.** Whole mount retinas from wildtype and *Kctd7*^−/−^ animals were stained for CD31 at P3, P8, and P12. In wildtype animals, the deep layer has not emerged at P3 but becomes visible by P8 and forms a complex and patterned plexus. In *Kctd7*^−/−^ animals, the deep layer is only sparsely present at P8 (**A**, scale bars = 250μm) and displays discontinuous vessel coverage and a reduction in growth across the retina (**B**, scale bars = 100μm). **C-D**. The defects in *Kctd7*^−/−^ deep layer vessel development were accompanied by a marked decrease in the area of the retina covered by the deep plexus (**C**) and a significant reduction in *Kctd7*^−/−^ deep layer vessel branch number (**D)** n=6 wildtype and 7 *Kctd7*^−/−^ animals. Data are represented as the mean ± the s.e.m. *** P<0.001, ** P<0.01, * P<0.05, 2way ANOVA and Dunnett’s multiple comparison test for significance.

### 3.3 Kctd7 may function in bipolar neurons to drive vasculature patterning

In the retina, each vascular layer can function as an individual neurovascular unit, such that changes to the vasculature can either induce or reflect local metabolic or neuronal defects (Paredes et al., 2018). We reasoned that while alterations to all layers of the vasculature might indicate that Kctd7 functions in the vasculature itself, the layer-specific defects we observed in *Kctd7*^−/−^ mice might indicate that Kctd7 could play a local or neuron-specific role. To begin to examine this, we performed *in situ* labeling over development to localize *Kctd7* expression. To examine key stages of this process, we first used probes against *Cspg4*, also known as *NG2* (a pericyte marker, Ozerdem et al., 2001), to visualize the vasculature (**Fig. 4**). As expected, *Cspg4* was detected in early development where it localized to thin bands. As development progressed, *Cspg4* labeling became more pronounced in the OPL, IPL, and GCL, consistent with the location of the deep, intermediate, and superficial vascular plexus. This pattern differed significantly from that of *Kctd7*. Unlike *Cspg4*, *Kctd7* was present in fate committed inner retina neurons as by P3 (**Fig. 4A**). From P8 to P12, *Kctd7* was present throughout the nuclear layers, with the highest levels of expression in the inner nuclear layer (**Fig. 4B-C)**. This pattern persisted into adulthood (**Fig. 4D**). Consistent with these results, *Kctd7* expression has been documented in retina neurons previously by both *in situ* and RNA-seq studies (Serial Analysis of Gene Expression, SAGE, https://cgap.nci.nih.gov/SAGE/mSEM Blackshaw et al., 2001; Blackshaw et al., 2004; Shekhar et al.,2016).

**Figure 4.**
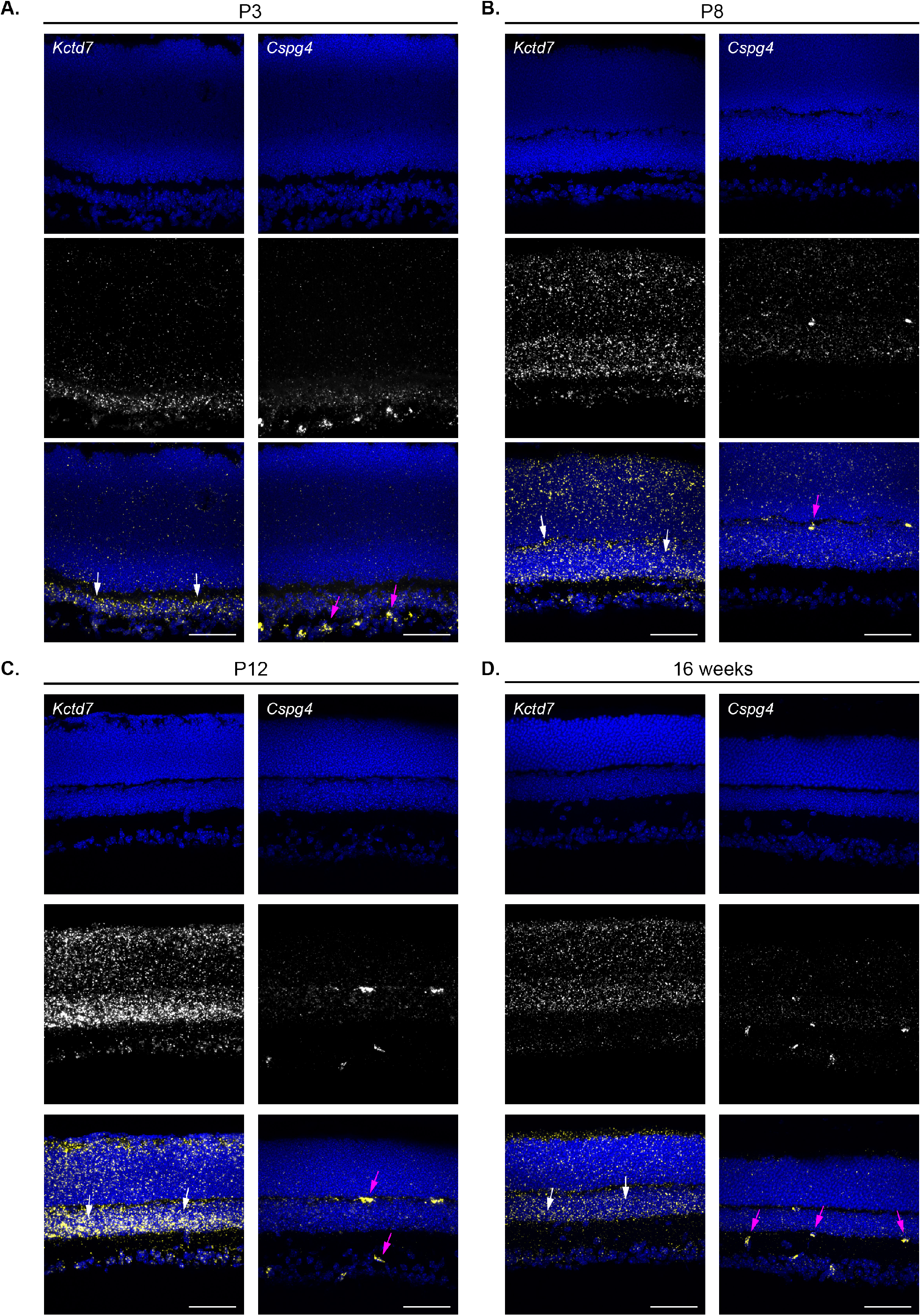
*Kctd7* is expressed in neurons and appears excluded from the vasculature. The presence and localization of *Kctd7* and the pericyte vasculature marker *Cspg4* were assayed in wildtype animals by florescent *in situ* hybridization at P3 (**A**), P8 (**B**), P12 (**C**), and 16 weeks (**D**). Consistent with the localization of retina vasculature, *Cspg4* was present in bands that colocalized with the retina synaptic layers (magenta arrows). In contrast, *Kctd7* was expressed broadly in retina neurons with an enrichment in the inner nuclear layer (white arrows) and did not display an expression pattern consistent with blood vessel localization. Scale bars = 50μm.

Because Kctd7 was expressed predominately in inner retina neurons, we next asked whether it may impact the organization or numbers of retina neuron subsets. To test this, we performed immunohistochemical assays to quantify each major retina neuron type at P8 when vascular defects emerge. We grouped the neuron markers based on the neurovascular unit that each cell type interfaces with. The superficial layer interacts with neurons in the GCL, which primarily includes RGCs (Sapieha et al., 2008; Edwards et al., 2012), while the intermediate and deep plexus are thought to interact with and support multiple retina neuron types that include bipolar cells, amacrine cells, horizontal cells, RCGs, and photoreceptors (Sun et al., 2015; Usui et al., 2015b, **Fig. 5A**). To examine these populations, we stained for markers that label each of these neuron types specifically (**Supplemental Table 1**). The numbers of RGCs, photoreceptors, amacrines, and horizontal cells were unchanged in *Kctd7*^−/−^ mice (**Fig. 5B-C**). Notably, however, bipolar cell development was altered. At P8 *Kctd7*^−/−^ mice displayed an 21% increase in the total number of bipolar cells relative to wildtype controls (1.47 ± 0.0.3 versus 1.23 ± 0.14 cells/500μm^2^, respectively, P<0.05, **Fig. 5D-E**). This increase appeared to be specific for the first postnatal week, as by P12 bipolar cell numbers were comparable in *Kctd7*^−/−^ and wildtype mice (1.26 ± 0.06 versus 1.30 ± 0.05 cells /500μm^2^, respectively), and numbers remained indistinguishable in adults (**Fig. 5E**). To determine whether increased cell death may account for the modulation of bipolar numbers over time, we co-stained for activated caspase-3 and the bipolar cell specific marker Chx10. Larger numbers of *Kctd7*^−/−^ bipolar cells were positive for activated caspase 3 during this period relative to wildtype controls (0.11 ± 0.04 versus 0.045 ± 0.02 cells /500μm^2^, respectively, P<0.01, **Fig. 5G-F**). These data suggest that Kctd7 alters bipolar cell numbers early in development and that these differences are normalized over time by a compensatory increase in bipolar cell death.

**Figure 5.**
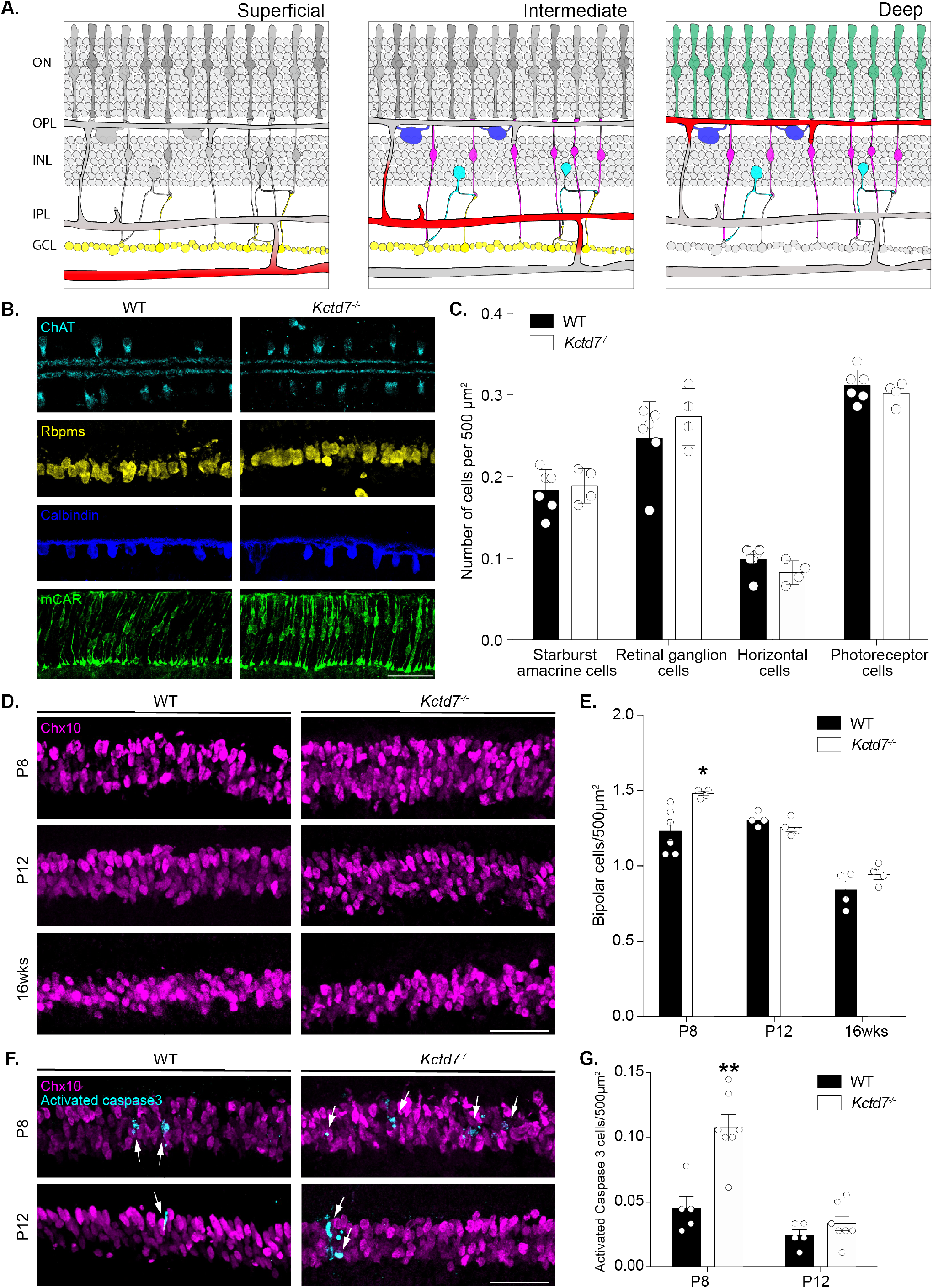
*Kctd7* mutant animals show an increased number of bipolar cell neurons. **A**. Representative schematics of the location of each major retina neuron type together with the vasculature network with which it is hypothesized to interact (yellow, RGCs; dark blue, horizontal cells; magenta, bipolars; cyan, amacrine cells; green, photoreceptors; red, blood vessels). **B**. Immunohistochemical images of each neuron type at P8 in wildtype and *Kctd7*^−/−^ mice: starburst amacrine cells (ChAT, cyan); retinal ganglion cells (Rbpms, yellow); horizontal cells (Calbindin, blue); cones (MCAR, green). **C**. Each neuron type was quantified at P8 in wildtype (n=5) and *Kctd7*^−/−^ mice (n=4) using antibodies specific for each population. No significant difference in the numbers of these neuron types were observed. **D-E.** Immunohistochemical images (**D**) and quantification (**E**) of bipolar cell nuclei following staining with Chx10 at P8, P12, and 16 weeks in wildtype and *Kctd7*^−/−^ mice. At P8, the number of bipolar cells was significantly increased (P=0.012), while bipolar cell numbers were not significantly altered at the other ages. **F-G.** The number of bipolar cells undergoing apoptosis was visualized (**F**) and quantified (**G**) in wildtype and *Kctd7*^−/−^ mice at P8 and P12 following co-staining for Chx10 and activated caspase 3. At P8, *Kctd7*^−/−^ bipolar cells showed a significant increase in the number of caspase positive nuclei relative to controls (P=0.0014, **G**). Scale bars = 50μm. Data are represented as the mean ± the s.e.m. *** P<0.001, ** P<0.01, * P<0.05, unpaired two-tailed Student’s *t* test.

### 3.4 Kctd7 is required for visual function

To determine the impact of Kctd7-mediated alterations on visual function, we performed electroretinography (ERG) on *Kctd7*^−/−^ mice to measure the integrity of photoreceptor responses (a-wave) and inner retina neuron responses (b-wave) under dark adapted and light adapted conditions. Significant reductions in both the a- and b-wave were observed in *Kctd7*^−/−^ mice under dark adapted conditions, resulting in an average 43% reduction of the a-wave and 34% reduction of the b-wave (**Fig. 6B-C**). Under light adapted conditions, the a-wave also trended lower (**Fig. 6D**), and significant reductions were observed in the b-wave (33% reduction at a flash intensity of 3 cd*s/m^2^, P=0.012, and a 36% reduction at a flash intensity of 10 cd*s/m^2^, P=0.03, **Fig. 6E**). Collectively, these results suggest that Kctd7-mediated changes in the retina disrupt visual function.

**Figure 6.**
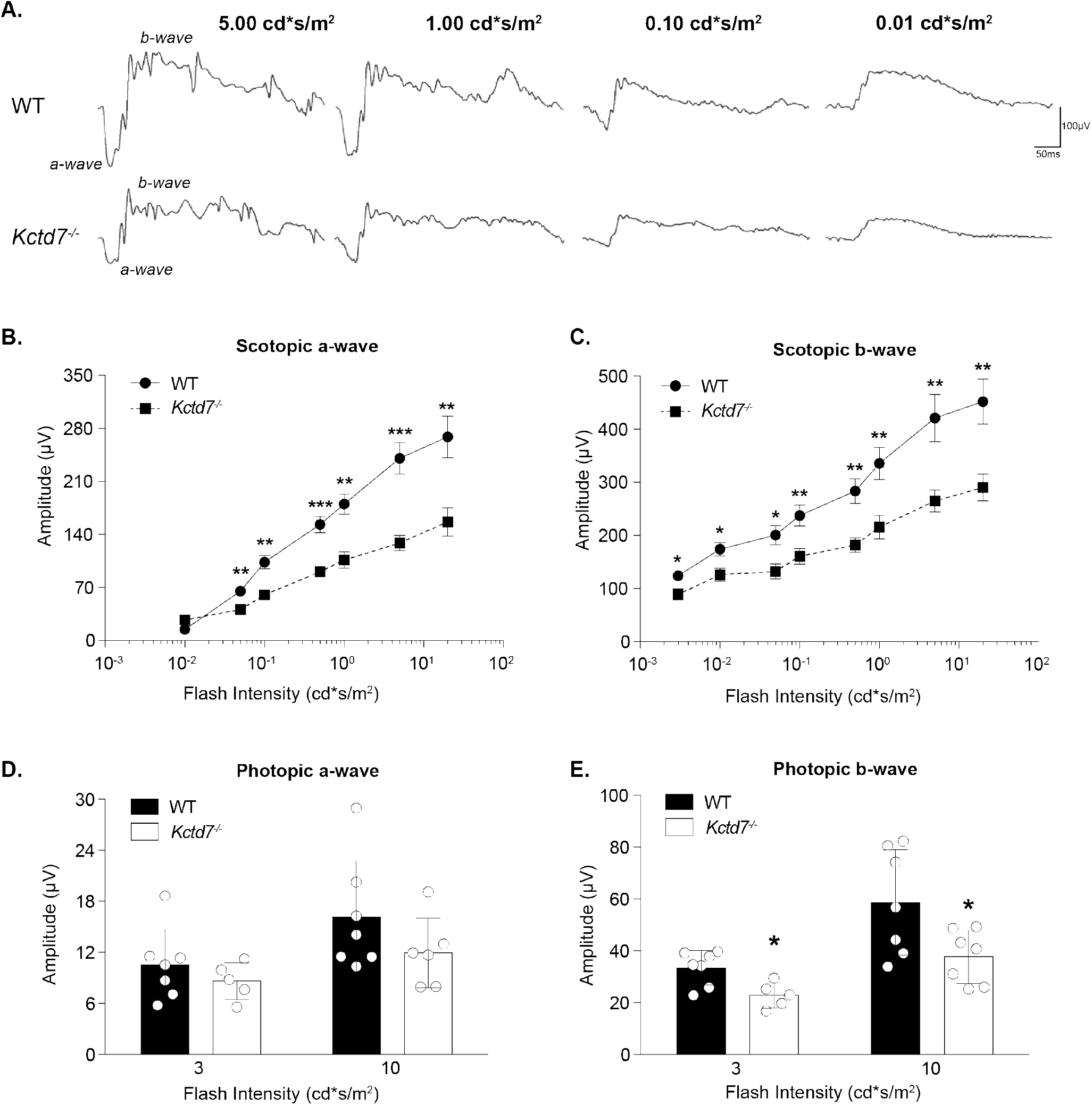
*Kctd7* is required for normal retinal function. Photopic and scotopic ERGs were recorded from 16-week-old wildtype (n=8) and Kctd7 mutant animals (n=7). **A**. Representative traces of 4 steps (0.01, 0.10, 1.00, and 5.00 cd*s/m^2^) are shown of ERG recordings under scotopic conditions. **B-C**. Under scotopic conditions, ERG recordings show a significant reduction in the a-wave amplitude for flash intensities greater than 0.003 cd*s/m^2^ (**B**) and in all b-wave amplitudes (**C**) relative to wildtype controls. **D-E**. Under photopic conditions ERG recordings trended lower for the a-wave but were not significantly different (**D**), while a significant reduction was observed in the b-wave at both 3.0 and 10.0 cd*s/m^2^ (**E**). Data are represented as the mean ± the s.e.m. *** P<0.001, ** P<0.01, * P<0.05, unpaired t-test using the Holm-Sidak method to correct for multiple comparison.

## 4. Discussion

The molecular programs that coordinate neuron development with vascular development have been challenging to resolve due to the heterogeneity and complexity of neurons and vessels in the brain. Here, we leverage the murine retina to examine the function of the progressive myoclonic epilepsy-associated gene *KCTD7*. Precise intraretinal ordering allows vessels to interact with and respond to neurons arrayed in the corresponding region of the retina (Stahl et al., 2010; Usui et al., 2015b; Selvam et al., 2017). In the absence of Kctd7, this process is defective: the deep and intermediate layers demonstrate increased vessel branching and vessel area in adults, and in development the emergence of both the superficial and intermediate layers is delayed. Thus, Kctd7 is required for vascular formation and organization. Kctd7 appears to act in part through cell-specific regulation of neuron survival, as Kctd7-deficient mice show a specific and time dependent developmental increase in the number of bipolar neurons that is corrected in adulthood. In keeping with these alterations, *Kctd7*^−/−^ mice displayed altered photopic and scotopic functional responses. Together, these results suggest a model in which Kctd7 acts as a conduit for neurovascular communication that impacts retinal function.

Three lines of evidence support a model in which Kctd7 influences vascular arrangement through its role in retina neurons. First, Kctd7 impacted only a subset of vascular layers rather than the entire network, suggesting that the changes are due to local differences in neuron form or function. In addition, the most affected layers were those closest to the neurons with high levels of *Kctd7*. Second, *Kctd7* was localized predominately to neurons. Indeed, *in situ* hybridization analysis showed that *Kctd7* is enriched in the INL where bipolar neurons reside. This neuron-restricted expression pattern is consistent with Kctd7 staining patterns in mouse brain, where it is enriched in specific neuron populations that include the olfactory bulb, deep layers of the cerebellar cortex, and Purkinje neurons (Azizieh et al. 2011). Third, *Kctd7* mutants showed increased numbers of bipolar neurons in the first postnatal week, while other neuron subsets were largely unaffected. Together, these results suggest that neuronal Kctd7 may help tune the precise timing and organization of intraretinal vascular development.

How might Kctd7 function in neurons to impact the vasculature? While the details of Kctd7 cellular biology remain obscure, data support potential mechanisms that center on potassium conductance (reviewed in Liu et al., 2013). KCTD7 is part of a family of proteins that contain the T1 tetramerization domain of voltage gated potassium channels (Kreusch et al., 1998; Stogios et al., 2005). In the inner retina of mouse and other species, potassium channels have been localized to bipolar cell dendrites where they form channels that rapidly activate after depolarization (Attwell et al., 1987; Kaneko et al., 1989; Karschin and Wässle, 1990; Klumpp et al., 1995). These outward currents can function to hyperpolarize the cell and restore membrane potential (Yazulla et al., 2001). Correspondingly, KCTD7 over-expression has been shown to hyperpolarize the cell membrane and reduce the excitability of transfected neurons in culture (Azizieh et al., 2011). Thus, it is possible that the alterations in vessel patterning we observe in *Kctd7*^−/−^ mice reflect neuron hyperexcitability. But how potential changes in potassium permeability occur is uncertain. In one proposed mechanism, Kctd7 may directly associate with GABAb receptor subunits and control potassium conductance through G protein-based regulation, as has been documented for KCTD8, 12, 12b, and 16 (Schwenk et al., 2010; Seddik et al., 2012; Fritzius et al., 2017). Alternatively, like other KCTD family members (Bayón et al, 2005; Smaldone et al., 2015; Ji et al., 2016; Pinkas et al., 2017), KCTD7 could bind the ubiquitin ligase CUL3 and induce the degradation of as yet unidentified substrate to modify potassium conductance (Azizieh et al., 2011). Since the levels of other potassium channels, including Kv1.2 and 1.5 and some members of the KCNQ family, can be regulated by ubiquitination, this is an interesting proposition (Henke et al., 2004; Boehmer et. al, 2008; Ekberg et al., 2007). Retina vasculature can also be regulated through CUL3-mediated ubiquitination of the Notch signaling pathway (Ohnuki et al. Blood, 2012), suggesting that Kctd7-CUL3 interactions also have the potential to participate in vessel remodeling directly.

Our ability to precisely follow the development of each vascular layer in *Kctd7* mutants allowed us to compare Kctd7-dependent phenotypes across time. We noted some interesting differences: the superficial and the deep layers failed to develop as quickly or completely as that in wildtype animals at P8, while in adults the intermediate and deep layers, but not the superficial layer were affected and instead showed increased vessel branching. What might be the causes of these differences? One potential explanation lies in the features of retina neuron development at P8, which represents a key time in retina maturation. First, it marks the end of new neuron generation (reviewed in Cepko, C., 2014). Second, bipolar cells have begun to form synapses with pre and post-synaptic partners (Onley et al. 1968; Blanks et al., 1974; Rich et al, 1997). Third, synaptic proteins like VGLUT1 become present, and synapses themselves begin to be active (Sherry et al., 2003). Thus, we speculate that during development Kctd7-dependent changes in neural activity may impact the integration of bipolar cells into developing synapses. Since retina vasculature can be highly structurally responsive to retina neural alterations (Sapieha et al., 2008; Fukushima et al., 2011; Edwards et al., 2012; Usui et al., 2015b), this imbalance may in turn influence not only bipolar cell development but also delay vessel emergence and complexity. How might these changes ultimately lead to increased vessel branching in adults? While more experiments will be needed to resolve this question, it is possible that like other bipolar cell potassium channels (reviewed in Yazulla et al.) Kctd7 may bring the membrane potential closer to the equilibrium potential after depolarization. Thus, it’s absence may ultimately result in adult neuron hyperexcitability leading to increased demand for nutrients and a corresponding increase in vessel branching. Alternatively, Kctd7 may have distinct functions at different developmental stages. For example, in the first postnatal week before synapses become active Kctd7 could influence bipolar cell survival and vessel emerge through non-activity dependent Cul3-mediated mechanisms. In contrast, it may impinge on adult neuron activity through its voltage gating properties. Finally, our data do not formally rule out a potential role for low levels of Kctd7 in the vasculature itself, and roles for potassium have been noted in vessel tone and blood flow (Filosa et al., 2006; Ladecola, C., 2017).

Our results join a small but growing field of related studies on other members of the KCTD family. This 26-member family shares sequence similarity with the voltage-gated K+ channel cytoplasmic domain, and these proteins consist of a conserved N terminus and a variable C terminus (Liu et al, 2013). While functions for the majority of these proteins are unknown, those that have been studied show a diverse array of properties that include cytoskeletal (Kang et al., 2004; Chen et al., 2009) and transcriptional regulation (Melnick et al. 2000; Amhed et al., 2003), as well as the ion channel gating and cullin E3 ubiquitin ligase functions discussed above. In addition, similar to KCTD7 mutations, several KCTD family members have been associated with human neural disorders (e.g. KCTD 12 and 13 [Golzio et al., 2012; Cathomas et al., 2015]), but the biological cause of these conditions has remained obscure in part because animal models have largely not been available. Since one of the criteria for IMPC generated lines is the absence of an existing null line, many Kctd family members have been entered into the IMPC pipeline (http://www.mousephenotype.org/), and several are undergoing analysis. While phenotype information for *Kctd7* has largely not yet been reported, data for several other family members are available (*Kctd1, 9, 10, 13, 17*) and suggest some interesting trends. All of the tested *Kctd* genes have at least one phenotypic alteration, and 3/6 have alterations to behavior or other neurologic functions (*Kctd9, 10, and 15*). In addition, 2/6 impact the heart or other vascular systems (*Kctd1 and 13*), and *Kctd1* also impacted the eye, showing persistence of the hyaloid vasculature and cataracts. Together with our findings, these data indicate that the KCTD protein family may play as yet underappreciated roles in nervous system development and function. In addition, these data may support screening human patients with mutations in KCTD7 or other related proteins for additional phenotypes that may impact health outcomes. Indeed, KCTD7-related human diseases have been sporadically associated with ocular phenotypes (Staropoli et al., 2012; Blumkin et al., 2012).

In summary, we have characterized a potassium channel tetramerization domain-containing protein responsible for coordinated emergence of the retina vasculature. We have also shown how an apparently neuron-specific gene can sculpt the organization of individual vascular layers, indicating new potential roles for Kctd family proteins. It is particularly attractive to consider this line for investigating the relationship between the numbers and activity of specific neuron types and layer-specific vessel arborization. Given the specific alterations in bipolar neurons in *Kctd7* mutants, further studies on the relationship between excitatory interneurons and vascular development may be warranted, particularly as they relate to progressive myoclonic epilepsy.

## Author Contributions

M.A.S. and J. A. designed the experiments. J.A. conducted the vessel and neuron imaging and analysis in adults and in development, performed the statistical analyses, conducted *in situ* and staining for Kctd7, and generated all of the figures. C.B. performed the neuron quantification and assisted with the *Kctd7 in situ*. D.J. performed the *in situ* for *Cpg4*. N.E.A. conducted the gene expression analyses and assisted with the figures. M.A.S. wrote the manuscript.

## Acknowledgements

We thank members of our laboratory and Jeffery L. Noebels for scientific discussions and advice. We also thank John R. Seavitt, Arthur L. Beaudet, Mary E. Dickinson, and their labs for help in the initial INSiGHT screen that led to this work and for providing the *Kctd7*^−/−^ mice. This work was supported by the National Institutes of Health (NIH, R00AG044444 and DP2EY02798 to M.A.S.), the Cancer Prevention Research Institute of Texas (RR150005), the Brain Research Foundation, and the Ted Nash Foundation. N.E.A. was supported by the National Institute of General Medical Sciences (T32GM088129). This project was also supported by the RNA *In Situ* Hybridization Core facility at Baylor College of Medicine with the assistance of Cecilia Ljungberg, Ph.D., and funding from the NIH (1S10 OD016167; IDDRC grant 1U54 HD083092, Eunice Kennedy Shriver National Institute of Child Health & Human Development). The availability of the *Kctd7* line was supported by KOMP2 awards UM1HG006348, U42OD11174, and U54HG006348.

## Competing Financial Interests

The authors declare no competing financial interests.

---

**Supplemental Figure 1.**
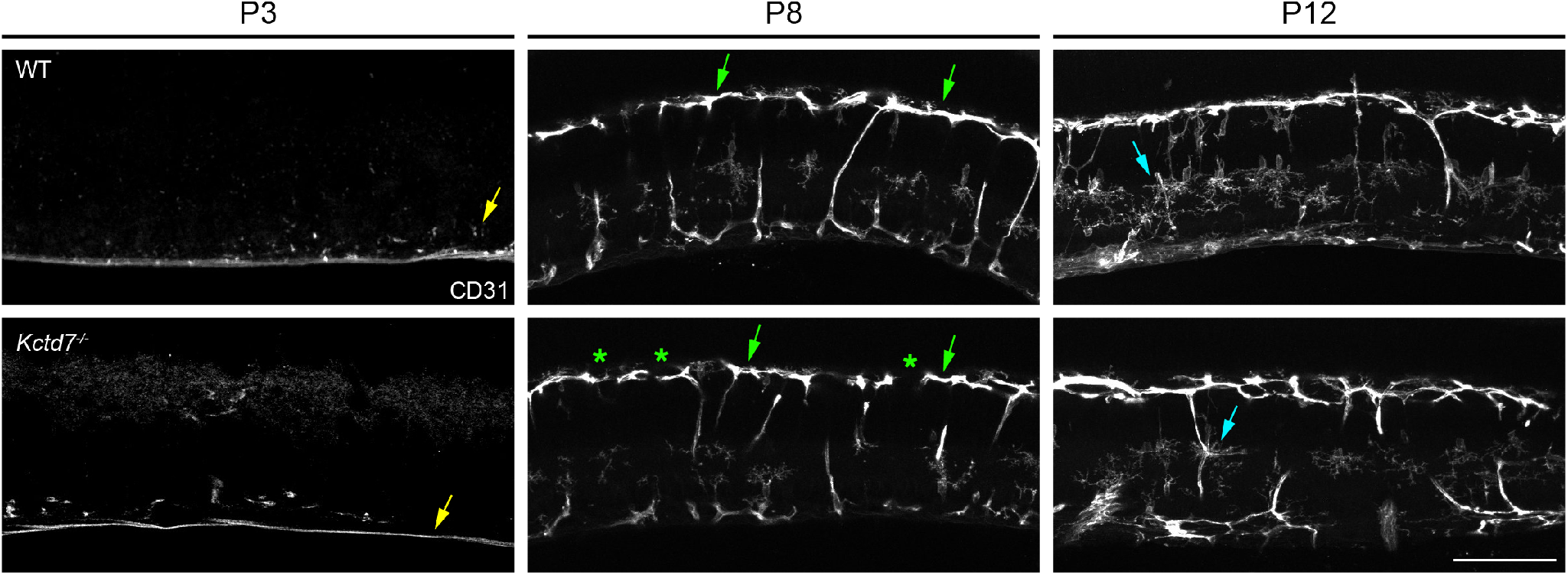
Laminar targeting of the vessel layers is intact in *Kctd7* mutant animals. Vessel laminar targeting was visualized in wildtype (upper panel) and *Kctd7*^−/−^ mice (lower panel) at P3, P8, and P12 following generation of retinal cross sections and staining for the vessel marker CD31. The superficial layer could be observed beginning at P3 (yellow arrows), the deep layer at P8 (green arrows), and intermediate layer at P12 (cyan arrows). Occasional gaps in the deep layer were present in *Kctd7*^−/−^ mice at P8 (green *), consistent with a reduction in deep vessel complexity at this time (see **Figure 3**). However, no large-scale defects in layer restriction were observed in *Kctd7* mutants. Scale bar = 100μm.

**Supplemental Table 1.** Antibodies used in *Kctd7* mutant tissue analysis. Antibodies were selected to reveal the vasculature and all major cell populations in the outer and inner retina.

| Antibody Name | Immunogen | Labelling Specificity | Source | Conc. |
| --- | --- | --- | --- | --- |
| Activated caspase-3 | Human Active Caspase-3 Fragment | Activated caspase-3 | BD Pharmingen; rabbit polyclonal; 559565: AB_397274 | 1:500 |
| Calbindin D-28k | Recombinant rat calbindin D-28k (CB) | Horizontal cells; amacrine cells; retinal ganglion cells | Swant; rabbit polyclonal; CB38; RRID:AB_10000340 | 1:10,000 |
| CD31 | 129/Sb Mouse-derived endothelioma cell line tEnd.1 | Platelet-endothelial cell adhesion | Fisher: rat monoclonal; BDB5500274; RRID:AB_393571 | 1:50 |
| Choline Acetyltransferase (ChAT) | Human placental enzyme | ChAT amacrine cells | Millipore; goat polyclonal; AB144P; RRID:AB_2079751 | 1:400 |
| Chx10 | Recombinant protein derived from the N terminal of the human Chx10 protein conjugated to KLH (aa 1 – 131) | Bipolar Cells | Exalpha; sheep polyclonal; X1180P; RRID:AB_2314191 | 1:300 |
| Cone Arrestin | Synthetic linear peptide | Cone photoreceptor | Millipore; rabbit polyclonal; AB15282; RRID:AB_11210270 | 1:5000 |
| Protein Kinase C alpha (PKCa) | Purified bovine brain protein kinase C | Rod bipolar cells | Abcam; mouse monoclonal; ab31; RRID:AB_303507 | 1:500 |
| Rbpms | Synthetic peptide corresponding to amino acid residues from the N-terminal region conjugated to KLH | Retinal ganglion cells | PhosphoSolutions; rabbit polyclonal; 1830-RBPMS; RRID: AB_2492225 | 1:250 |
| Syntaxin-1 | Synaptosomal plasma-membrane fraction from adult rat hippocampus | Pan-amacrine cells; horizontal cells | Sigma-Aldrich; mouse monoclonal; S0664; RRID:AB_477483 | 1:500 |
| Vglut3 | Recombinant protein corresponding to AA 543 to 601 from mouse VGLUT3 | Vglut3-expressing amacrine cells | Synaptic System; 135204; guinea pig polyclonal; RRID: AB_2619825 | 1:2000 |

**Supplemental Table 2:**
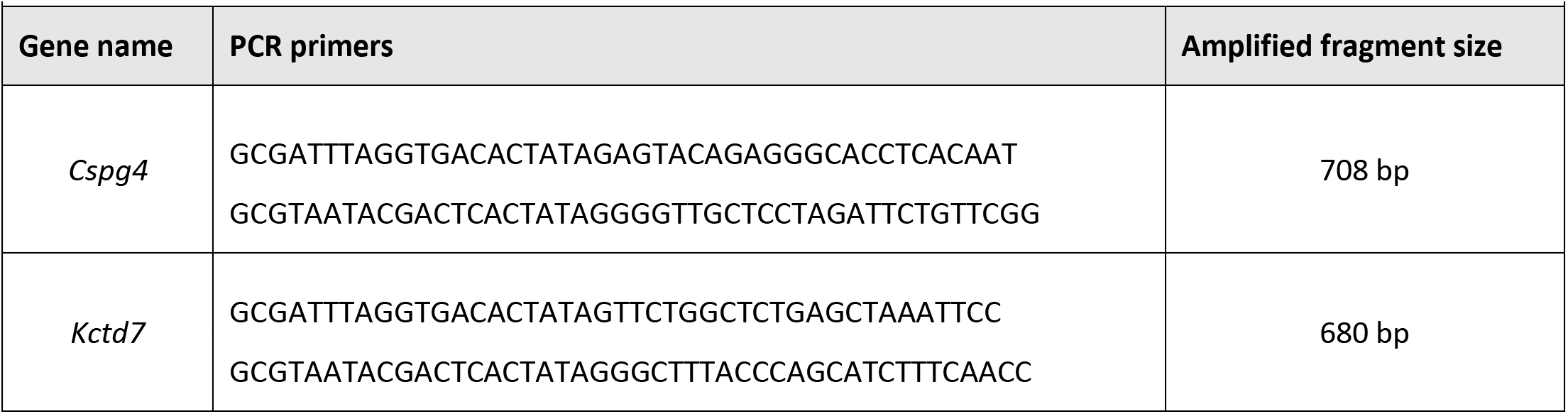
PCR primers used to amplify *in situ* probes.

